# Approximate Bayesian Computation Algorithms for Estimating Network Model Parameters

**DOI:** 10.1101/106450

**Authors:** Ye Zheng, Stéphane Aris-Brosou

## Abstract

Studies on Approximate Bayesian Computation (ABC) replacing the intractable likelihood function in evaluation of the posterior distribution have been developed for several years. However, their field of application has to date essentially been limited to inference in population genetics. Here, we propose to extend this approach to estimating the structure of transmission networks of viruses in human populations. In particular, we are interested in estimating the transmission parameters under four very general network structures: random, Watts-Strogatz, Barabasi-Albert and an extension that incorporates aging. Estimation was evaluated under three approaches, based on ABC, ABC-Markov chain Monte Carlo (ABC-MCMC) and ABC-Sequential Monte Carlo (ABC-SMC) samplers. We show that ABC-SMC samplers outperform both ABC and ABC-MCMC, achieving high accuracy and low variance in simulations. This approach paves the way to estimating parameters of real transmission networks of transmissible diseases.

## 1. Introduction

The essential idea behind Bayes’ theorem can be shown in the following equation is to relate the posterior distribution of parameter *θ* given the data *D*, or *P* (*θ|D*), to the likelihood of the data, or *P* (*D|θ*), as:

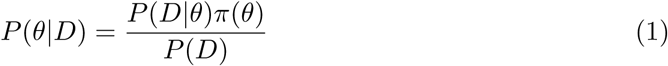

where *π*(*θ*) in Equation (1) is the prior distribution. The denominator, *P*(*D*) can be calculated as:

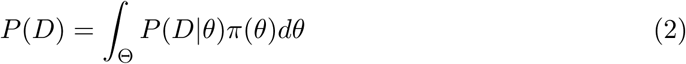

which is a constant that is extremely difficult to compute. In most applications, we may be interested in the posterior distribution. Therefore, once we are provided with the likelihood function and the prior distribution, the changing pattern of the parameters of interest can be obtained by *f* (*θ|D*) ∝ *f*(*D|θ*)*π*(*θ*), where *f* (.) potentially represents a density.

However, when the likelihood is expensive to compute or even, in some extreme case, unknown or extremely complicated as with transmission networks (Bickel and Chen, 2009), resorting to Approximate Bayesian Computation (ABC; Pritchard et al, 1999; Beaumont et al, 2002) is required.

To simplify the problem, we here consider four standard network architectures: Erdös-Rényi (ER; Erdös and Rényi, 1959) or random graphs, Watts-Strogatz (WS; Watts and Strogatz, 1998) graph, Barabasi-Albert (BA; Barabasi and Albert, 1999) graph and Aging and Preferential attachment (AP; Csardi and Nepusz, 2006). We implement parameter estimation under these four network architectures under the ABC framework and evaluate their performance in terms of accuracy and variance.

## 2. Materials and Methods

### 2.1 Approximate Bayesian Computation

The essential idea behind ABC is simulating data sets *D*^⋆^ from the sampled parameter *θ*^⋆^ following the prior distribution *π*(*θ*). Then the simulated the data sets *D*^⋆^ that are simulated are compared with real one *D*. If the simulated data sets *D*^⋆^ are close enough to *D*, then the parameter *θ*^⋆^ sampled from the prior distribution *π*(*θ*) is close to the real parameter *θ*. In this case, the sampled parameter *θ*^⋆^ is “accepted”, otherwise the simulation is “rejected”. The basic ABC algorithm is then summarized as:

1. Sample *θ*^⋆^ ∼ *π*(*θ*)
2. Simulate data set *D*^⋆^ ∼ *f* (*D*|*θ*^⋆^)
3. Compute the distance *d*(*D*^⋆^, *D*)
4. If *d*(*D*^⋆^, *D*) ≤ *ϵ*, accept *θ*^⋆^; return to Step 1.

The final estimation of the parameter *θ*, is then based on the posterior *P* (*θ|D*), here approximated by *P* (*θ|d*(*D*^⋆^, *D*) *≤ ϵ*).

There are basically two problems left in this basic algorithm. First is how to compare the simulated data set with the observed one so that the distance between them can be computed. The most common way is to adopt summary statistics *S* abstracting information from the data set *D* and calculate the distance *d*(*S, S*^⋆^) between these two summary statistics vectors. Inevitably, the information in the data set will shrink once it is represented by the summary statistics, as these are almost always inconsistent. However, the ABC sampler can correct for this with postsampling regression adjustment (Beaumont et al, 2002; Toni et al, 2009; Leuenberger and Wegmann, 2010). Furthermore, Euclidean distance or Mahalanobis distance can be employed to evaluate the discrepancy and the choice depends on the specific situation. The second potential problem is about defining the threshold *ϵ*: the smaller *ϵ* the closer simulations are to reality, but the lower the acceptance rate of the rejection sampler (ABC). Here, we offer two statistical approaches to determine *ϵ*.

#### 2.1.1 Determination of ϵ with Large Numbers of Iterations

By repeating the same simulation for very large numbers of time, we can obtain the empirical distribution of the distance variable defined above. If the iteration is large enough, the empirical distribution can be regarded as the real distance distribution. Then, for a given acceptance rate, it is easy to obtain the corresponding threshold value *ϵ*. For example, if the targeted acceptance rate is 1%, the 1% confidence interval under the empirical density curve around the observed summary statistics will be the acceptance area while the rest is the rejection area. In this case, we can guarantee that the final acceptance rate is approximately 1%.

#### 2.1.2 Determination of ϵ with Bootstrap Method

Bootstrap is resampling the data elements with replacement from the original data set to do point or interval estimation (Efron and Tibshirani, 1993). Therefore, we can resample a given data set *D* with replacement to form a group of similar data sets. Consequently, the confidence interval of the original data can be calculated, and hence the threshold *ϵ* can be defined as the critical value depending on a fixed significant level (*e.g.*, 5%).

While *ϵ* can be determined easily, in practice however chances are that the prior distribution is far from the posterior, so that acceptance rates are low and parameter estimation imprecise (Beaumont et al, 2002; Toni et al, 2009; Leuenberger and Wegmann, 2010). To improve accuracy of the naïve ABC sampler and reduce estimation bias, two directions are explored next. One is to adjust the estimation process by adding relative weights, while the other is to upgrade the ABC algorithm.

### 2.2 Postsampling Regression Adjustment and Weighting

The ABC sampler is based on whether the difference between simulated data set *D*^⋆^ and the observed data set *D* is small “enough”. Nevertheless, after accepting suitable samples, there still exists a discrepancy, *|d*(*D, D*^⋆^)*|* or *|d*(*S, S*^⋆^)*|* that should be eliminated. Therefore, local regression was suggested to reduce this discrepancy (Beaumont et al, 2002; Csilléry et al, 2010; Leuenberger and Wegmann, 2010). The most straight-forward approach is to use Least Square (LS) to fit the accepted samples to reduce bias as:

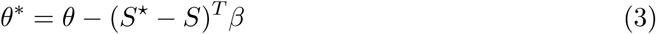

where *β* is the local linear regression slope. More sophisticated, non-linear regression models, have been proposed (Blum and François, 2010) (non-linear regression models for approximate bayesian) but we show below that even the simple LS regression does not always correct bias.

### 2.3 ABC-MCMC

The combination of the naïve ABC algorithm described above with Markov chain Monte Carlo (MCMC) method leads to ABC-MCMC samplers (Marjoram et al, 2003). These can be described as:

1. Initialize *θ*_*i*_ (*i* = 1)
2. Propose a candidate sample *θ*^⋆^ from *θ*_*i*_ following the transition distribution *θ*^∗^ ∼ q(*θ|θ*_*i*_)
3. Simulate data set *D*^⋆^ ∼ *f* (*D|θ*^⋆^)
4. If *d*(*S*(*D*^⋆^), *S*(*D*)) *≤ ϵ*, go to 5, otherwise return to 2
5. Set *u ∼* Uniform[0, 1], if 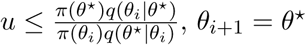, otherwise *θ*_*i*+1_ = *θ*_*i*_
6. Return to 2.

This algorithm was shown to converge on the target distribution *f* (*θ|d*(*S*(*D*^⋆^), *S*(*D*)) *≤ ϵ*) (Marjoram et al, 2003).

While the ABC-MCMC algorithm has larger acceptance rates than the naïve ABC method, fine-tuning the sampler can be difficult (Haario et al, 2001).

### 2.4 ABC-SMC

To improve on ABC-MCMC samplers, Sisson et al (2007) proposed the ABC Sequential Monte Carlo (ABC-SMC) sampler, which can be seen as an extension of importance sampling (Liu, 2001).

The idea underlying the algorithm is to generate several populations or sampling levels, simulating parameter values from their prior distribution, and then resampling from the previous samples (from the previous level). This process is repeated for a pre-defined number of levels. At each level, samples for which the distance between the simulated data set *D*^⋆^ and the observed data *D* is small enough will remain in the population, otherwise they will die out through more and more strict selection process. The particles are propagated and selected through a sequence of intermediate distributions *π*(*θ|d*(*D, D*^⋆^) *≤ ϵ*) (Toni et al, 2009). Moreover, the selection criterion, which is a set of threshold values *E*, is becoming more stringent: *ϵ*_1_ > *ϵ*_2_ >… > *ϵ*_*T*_ ≥ 0. By restricting this evolution process from the prior to the target distribution, the potential disparity in simulating the parameters is avoided (Sisson et al, 2007).

Algorithm of ABC-SMC can then be summarized as (Toni et al, 2009):

1. Set the threshold values *ϵ*_1_, *…, ϵ*_*T*_ together with population indicator t = 0
2. Set particle indicator *i* = 1 and go through the Steps in ABC to get the first population sample and denote the weight for each particle as 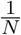. When *t* > 0, sample *θ*^⋆^ from the previous population {*θ*_*t−*1_} with weights {*w*_*t−*1_} and perturb the *θ*^⋆^ to get *θ*^∗^ ∼ *q*(*θ*|*θ*^⋆^)
3. Simulate data set *D*^⋆^ *∼ f* (*D|θ*^*∗*^). If *d*(*S*(*D*^⋆^), *S*(*D*)) *≤* ϵ, go to 4, otherwise return to 2
4. Set *θ*_*t*_ = *θ*^*∗*^ and calculate the corresponding weights 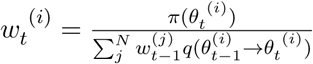
5. Normalize the weights. If *t < T*, set t = t + 1, return to 2.

Unlike ABC-MCMC, ABC-SMC is neither stuck in a low likelihood area nor generate dependent particles (Sisson et al, 2007). Also, ABC-SMC does not need to require any burn-in period. However, this algorithm spends most of its time constructing the auxiliary distributions which are not directly used in the final estimation (Sisson et al, 2007). Additionally, there is no general rule for setting the sequence value of the threshold (*ϵ*).

## 3. Results

In this section we present simulation results to assess accuracy, as measured by mean squared errors (MSEs) and variance of network estimation under the three samplers, ABC, ABC-MCMC and ABC-SMC. These simulations are based on networks containing 170 nodes and are made of 50 replicates. Parameter estimation is performed under the four networks described above: ER has one parameter (*α*: the connection probability), WS has two (*α*: rewiring probability; *β*: neighborhood size), BA has one (*α*: preferential attachment) and AP has two (*α*: preferential attachment; *β*: aging).

### 3.1 ABC sampler

In the naïve ABC algorithm, we have introduced in detail two main algorithms, these are the rejection sampler and the adjust rejection sampler. Usually, after adjustment, the density curves tend to have more samples gathering around the true parameter consequently resulting in a sharp peak by the true parameter value. However, the adjustment has distinct effect on the four network model as is shown in Figure 1. While estimation is improved dramatically for the ER model and to some extent for WS model, the BA and AP models show no improvement to slightly worse estimates.

When simulated under different parameter values, we can observe that local linear adjustment has a positive impact on parameter estimation for ER and WS under any value of *α*, while for BA and AP the MSE can be worsen after weighting (Figure 2).

**Figure 1:**
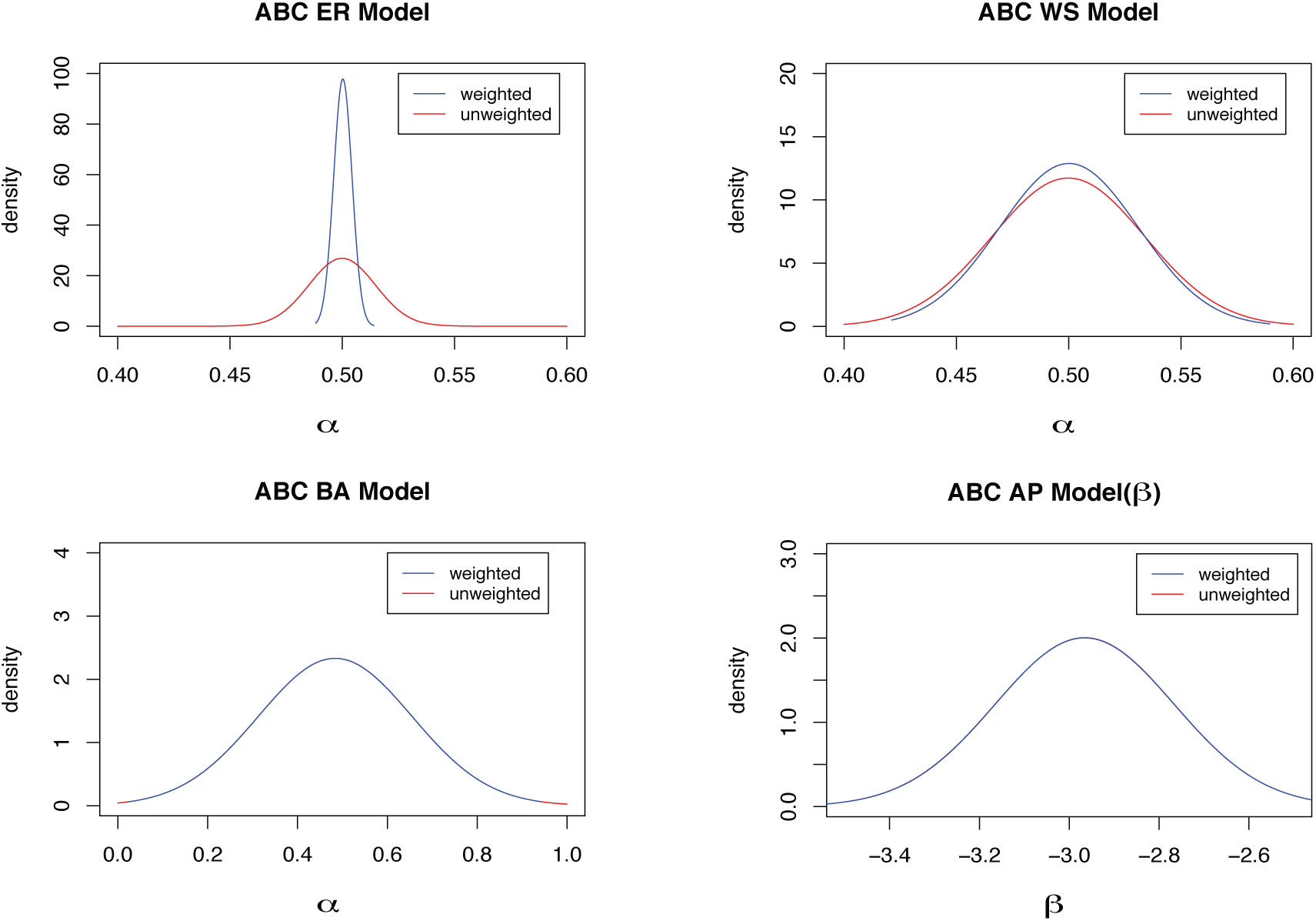
Estimated density distributions under ABC. Red curves represent unadjusted samples, while the blue ones are for adjusted results. The true parameter values for the four models are 0.5 for ER (*α*), WS (*α*) and BA (*α*) while it is -3 for AP (*β*).

For BA model, if the true parameter value is approximately *≥* 2, MSE increases after weighting. This bias is due to a positive or negative slope of the linear local adjustment that, depending on the direction of the shift, will lead to a larger deviation (BA curve in Figure 2). For AP, the observed bias is the result of fixing *β* to estimate *α* and *vice versa*. Due to the large bias and uncontrolled instability, the *alpha* parameter cannot be reliably estimated with this ABC sampler.

A similar results can be seen in Figure 3, where *α* for AP is unstable most of time and the estimation error is relatively large compared to *β* in the same model as well as the rest three models. Therefore, if applied to real data set, the estimation of *α* under AP model will be unconvincing. Fortunately, BA model, which merely has one parameter, is a simpler model of AP whose coefficient and constant term of *β* is zero. In other word, the BA model can, to some extent, demonstrate the behavior of *α* in AP model.

**Figure 2:**
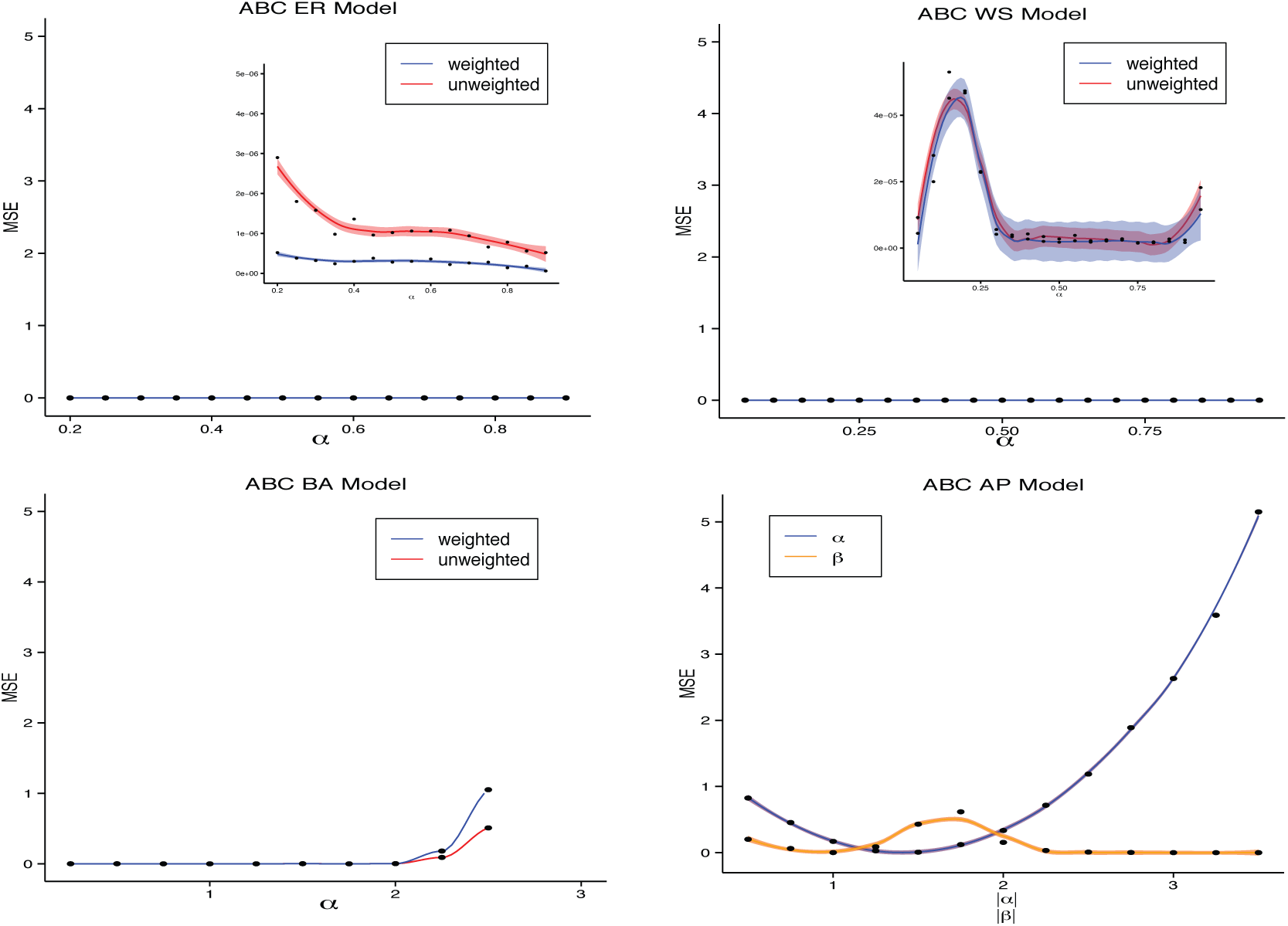
MSE results for ER, WS, BA and AP. The mode of the density estimated by ABC is used for estimation.

WS has two parameters, just like AP. However, estimation is more accurate and stable for WS model than for AP model. Generally speaking, the variance and MSE is low enough to guarantee that stability and accuracy will not change dramatically for any fixed neighborhood size (Figure 4). More specifically, both the variance and MSE have a tendency to decrease with increasing neighborhood sizes (insets in Figure 4). From Figure 4, it can be concluded that neighborhoods larger than 40, the variance and especially the MSE become extremely small. Therefore, without including all the vertices of the network, we can still achieve a near-perfect estimation.

### 3.2 ABC-MCMC

We may start from the result simulated by ABC-MCMC method on WS model (Figure 5).

While ABC-MCMC can achieve a reasonable accuracy and stability, the choice of variance for the proposal kernel can impact mixing. In spite of the accurate estimation, the sampler can waste a long time before sampling from the target distribution, starting from a random initial point (Figure 6).

**Figure 3:**
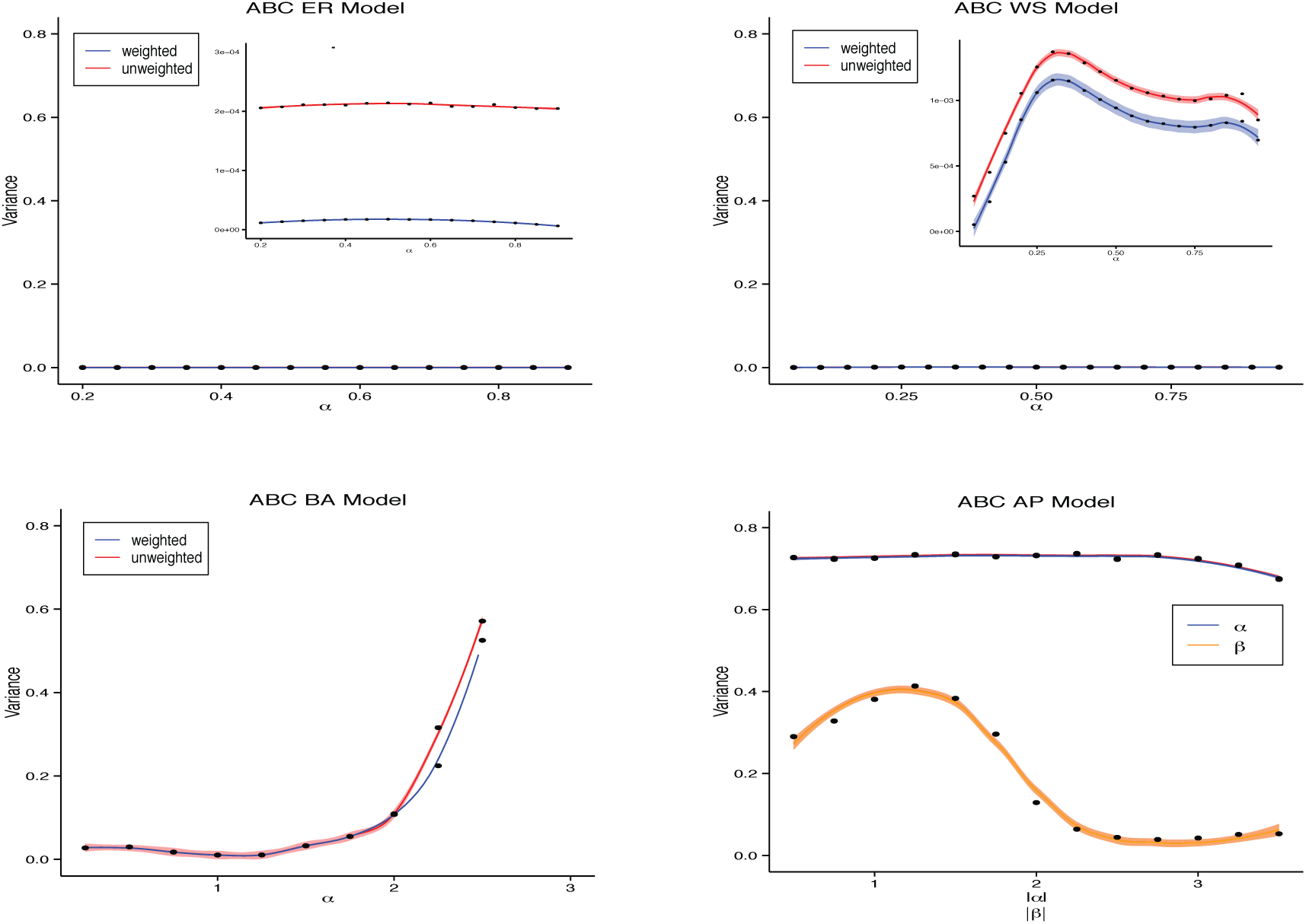
The variance of the accepted sampler in estimating the true parameter. It is obvious to figure out that variances in ER model and WS model have different magnitude partially due to the distinguished range of the parameters in each of the network model.

### 3.3 ABC-SMC

Under ABC-SMC, the long period of waiting time for the Markov chain under ABC-MCMC to forget its initial state does not happen. Moreover, Figures 7 and 8 show that both MSE and the variance are much smaller than under naïve ABC.

We also provide the empirical density distribution directly showing the evolving process of SMC in Figure 9.

**Figure 4:**
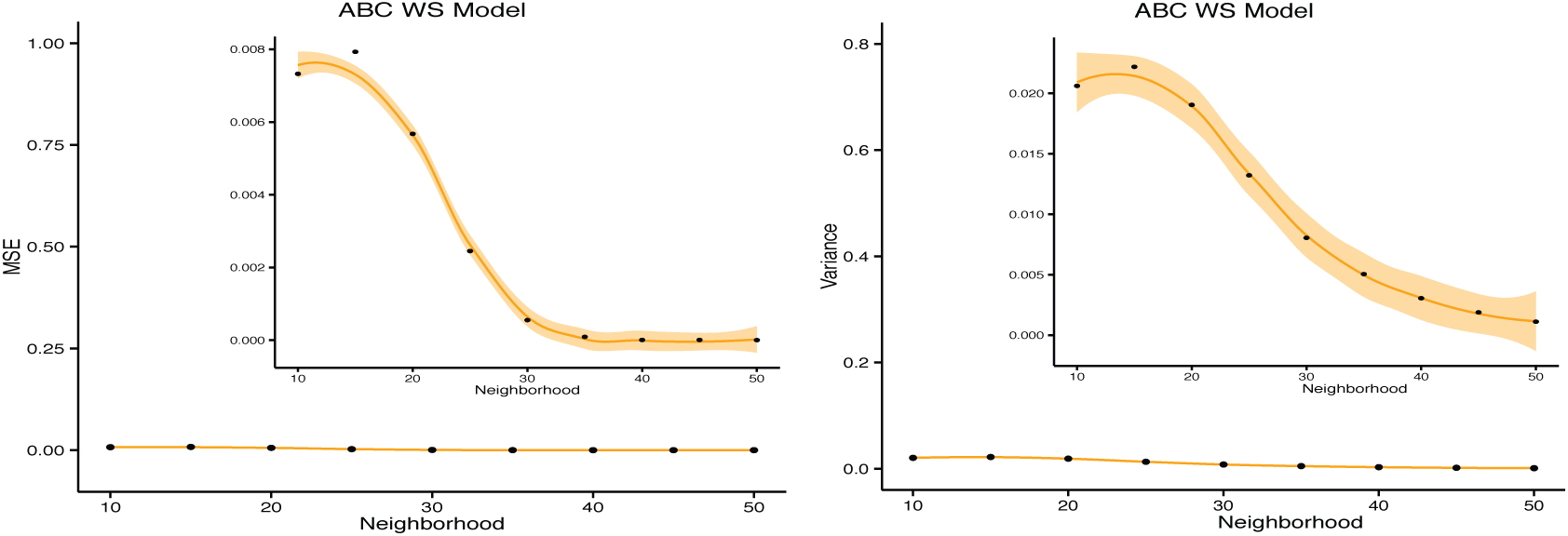
MSE and variance for WS under different neighborhood value. WS model has two parameters which are rewiring probability and the neighborhood. Neighborhood here means within how many neighbor distance the vertices of the lattice will be connected. Rewiring probability fixed at 0.5, we explore the changing pattern for different neighbor size in the final estimation. The shaded area represent 95% confidence interval of the smoothing kernel.

## 4. Discussion

In this paper, we have discussed about the basic ABC algorithm and the improved versions with ABC-MCMC and ABC-SMC. We showed by simulations that application of ABC methods to estimate the network parameters is accurate and stable. Estimation under the naïve ABC can be biased and unstable in estimating network parameters, while ABC-MCMC is sensitive to both initial values and proposal kernel tuning, which need to be set before running ABC-MCMC algorithm (but see adaptive samplers: Haario et al, 2001). Consistently with others (Sisson et al, 2007), our results show that ABC-SMC achieves both low variance and high accuracy. Therefore, in analyzing the real data on transmission networks, ABC-SMC is expected to perform better.

**Figure 5:**
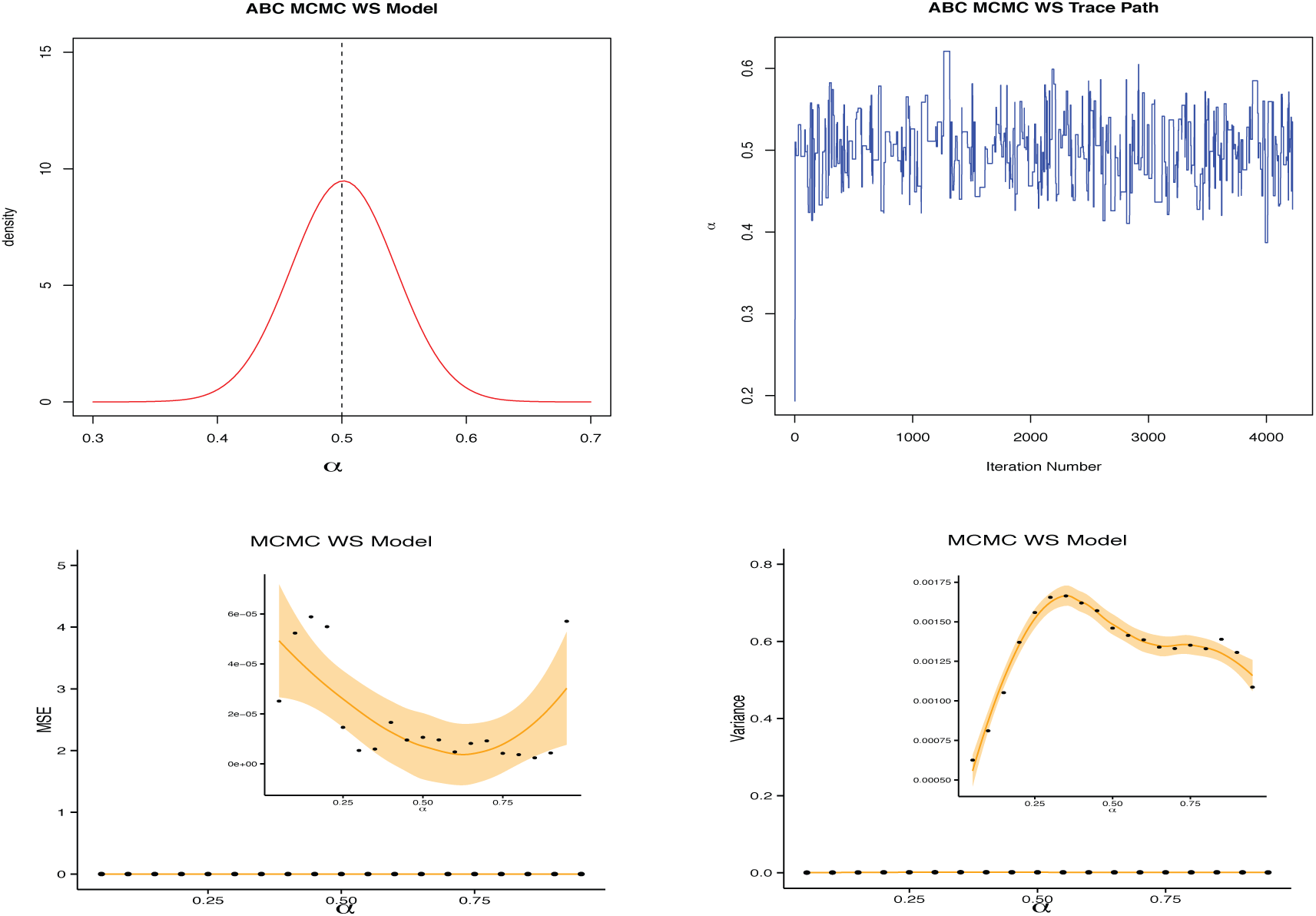
Thorough description of the ABC-MCMC algorithm. WS is used to illustrate. Vertical dashed line: true parameter value.

**Figure 6:**
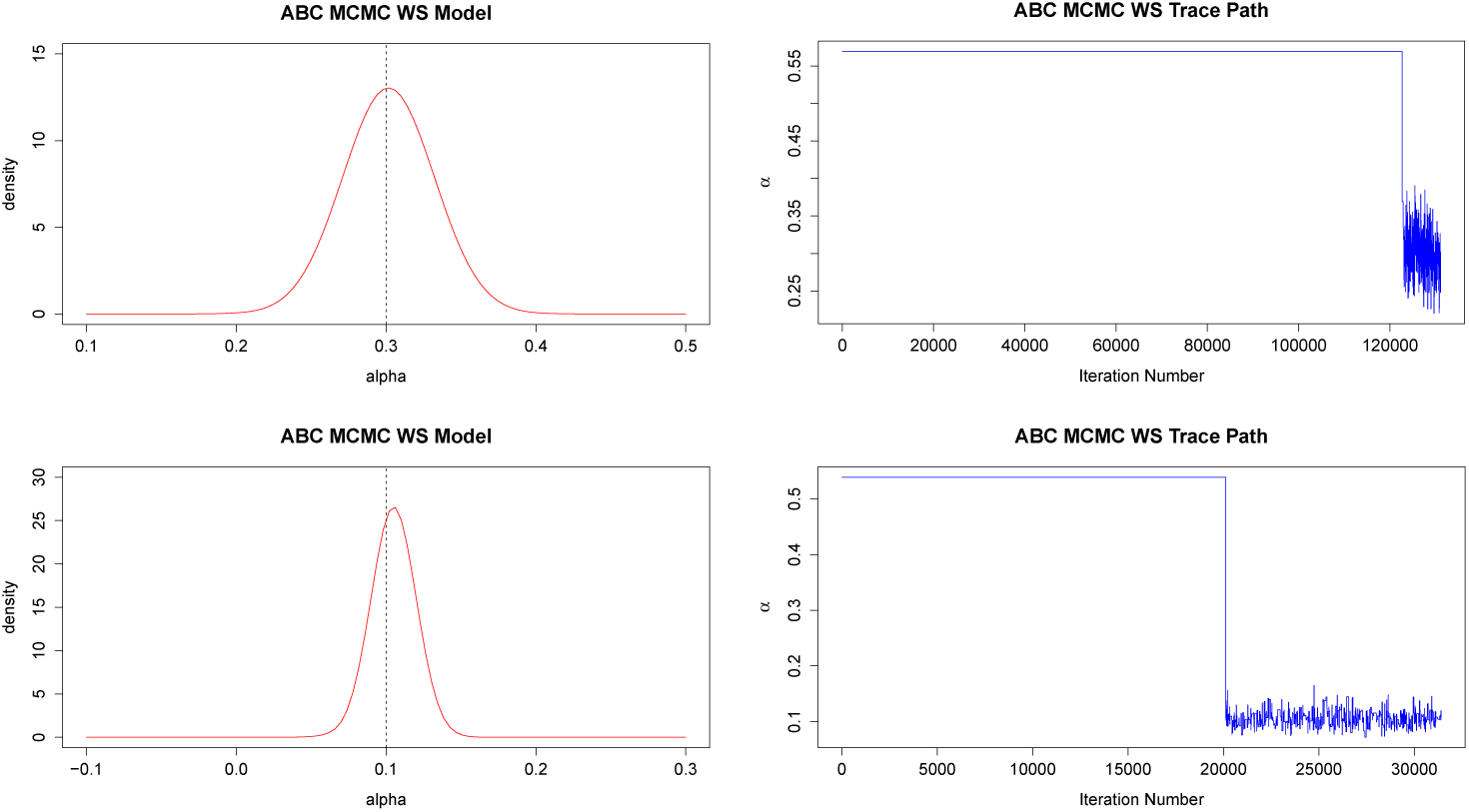
Estimation and trace path under ABC-MCMC. The empirical density distribution is an approximate symmetric distribution and the highest peak, the mode, is defined as the estimation for the true parameter. Dash line represents the true parameter value.

**Figure 7:**
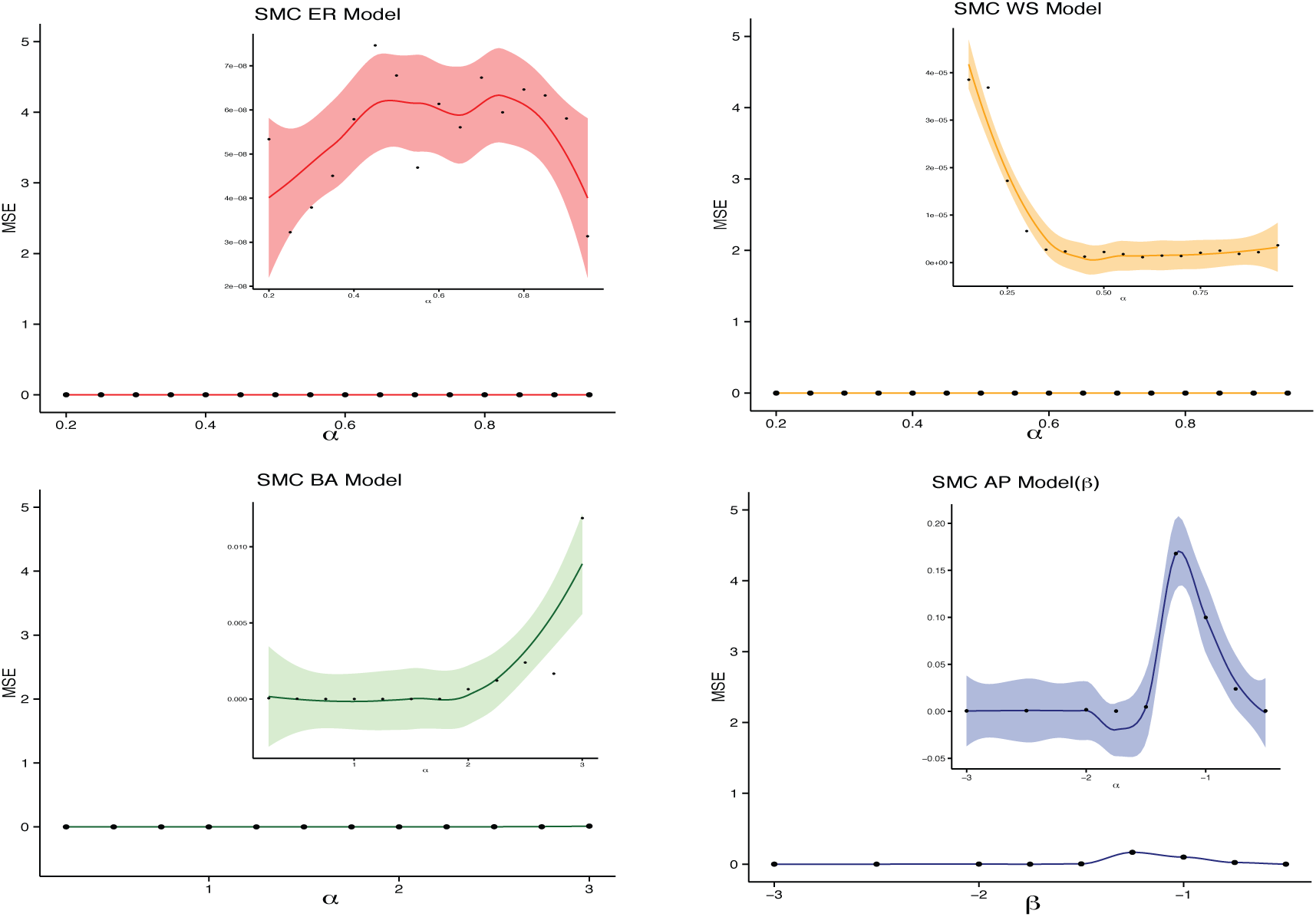
MSE under ABC-SMC for ER, WS, BA and AP model. Shaded areas represent 95% confidence intervals for the smoothing kernels.

**Figure 8:**
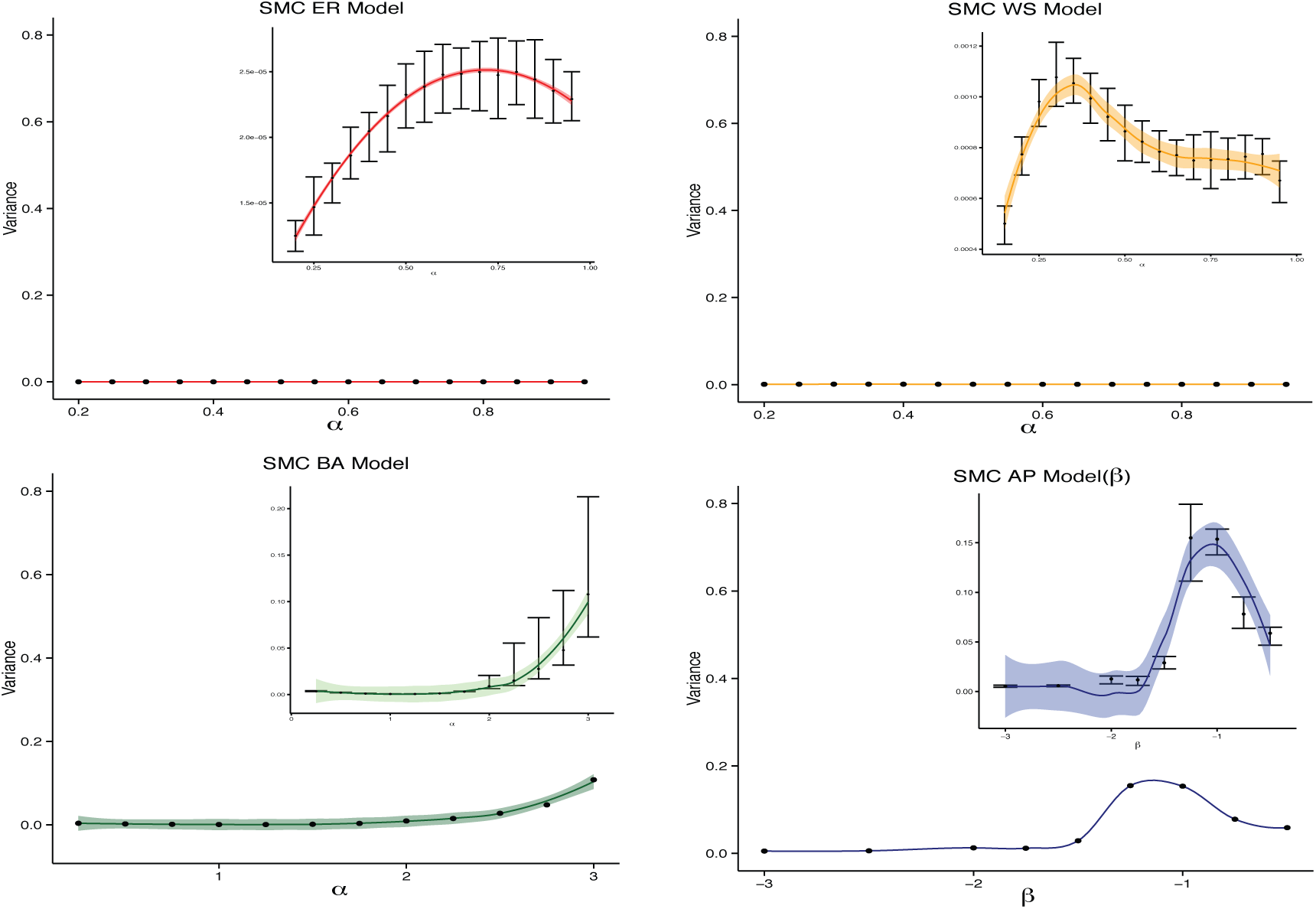
Variance under ABC-SMC for ER, WS, BA and AP model. Error bars represent empirical 90% confidence intervals, while shaded areas represent 95% confidence intervals for the smoothing kernels.

**Figure 9:**
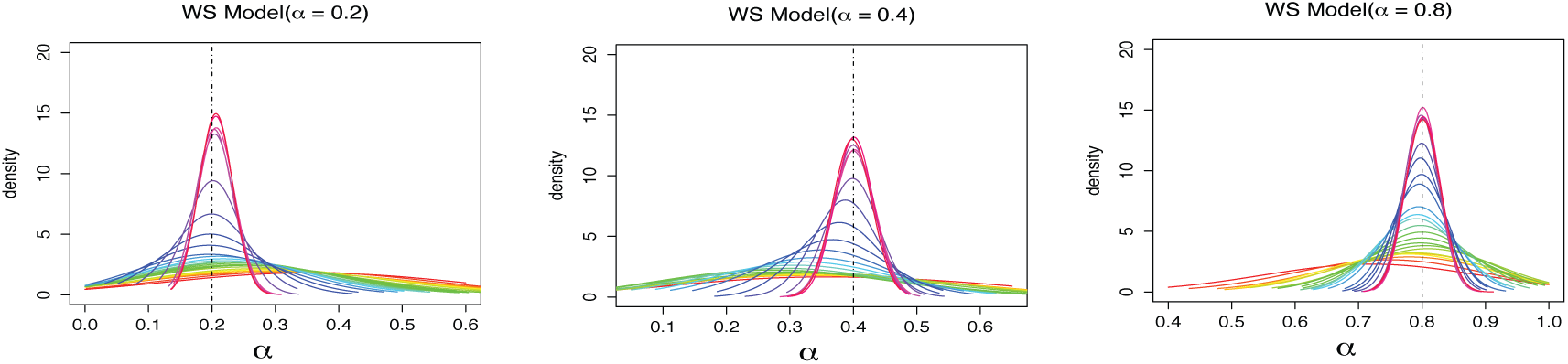
The empirical density distribution for each evolving step under its corresponding threshold for WS. Similar results obtained under other networks (not shown).

## Acknowledgements

This work was supported by MITACS Globalink and the Chinese Scholarship Council (YZ) and the Natural Sciences and Engineering Research Council of Canada (SAB).

